# Genomics to Notebook (g2nb): extending the electronic notebook to address the challenges of bioinformatics analysis

**DOI:** 10.1101/2023.04.04.535621

**Authors:** Michael Reich, Thorin Tabor, John Liefeld, Jayadev Joshi, Forrest Kim, Helga Thorvaldsdottir, Daniel Blankenberg, Jill P. Mesirov

**Affiliations:** Department of Medicine, School of Medicine, University of California, San Diego, La Jolla, CA, USA; The Broad Institute of MIT and Harvard, Cambridge, MA, USA; Moores Cancer Center, University of California, San Diego, La Jolla, CA, USA; Genomic Medicine Institute, Lerner Research Institute, Cleveland Clinic, Cleveland, OH, USA; Department of Molecular Medicine, Cleveland Clinic Lerner College of Medicine, Case Western Reserve University, Cleveland, OH, USA

## Abstract

We present Genomics to Notebook (g2nb), an environment that combines the JupyterLab notebook system with widely-used bioinformatics platforms. Galaxy, GenePattern, and the JavaScript versions of IGV and Cytoscape are currently available within g2nb. The analyses and visualizations within those platforms are presented as cells in a notebook, making thousands of genomics methods available within the notebook metaphor and allowing notebooks to contain workflows utilizing multiple software packages on remote servers, all without the need for programming. The g2nb environment is, to our knowledge, the only notebook-based system that incorporates multiple bioinformatics analysis platforms into a notebook interface.

---

The electronic notebook has become the de facto medium for much of data science and bioinformatics due to its accessibility and ease in incorporating scientific exposition with executable code to form a reproducible “research narrative.” Jupyter Notebook[1] and its successor JupyterLab[2] are the preeminent platforms. However, there are well-established bioinformatics analysis platforms, such as Galaxy[3] and GenePattern[4], that collectively host thousands of tools on their own compute resources. A system that could incorporate these platforms into the notebook metaphor would bring the benefits of their tools to notebook users while avoiding the installation, compatibility, and resource requirement issues that often arise when notebook developers add external packages to a Juypter environment. To address these issues, we have released Genomics to Notebook (g2nb), which builds on JupyterLab to add access to multiple bioinformatics platforms and other functionality through the components described below. While we focus on functionality for the non-programmer, we note that all programmatic features of JupyterLab are retained, making g2nb appealing to a wide range of users.

The g2nb environment incorporates multiple bioinformatics software platforms within the notebook interface. A standard Jupyter notebook consists of a sequence of cells, each of which can contain text or executable code. We have added a new cell type that provides an interface within the notebook to tools that are hosted on a remote Galaxy or GenePattern server. These new *analysis* cells present a form-like interface similar to the web interface of the original platforms, requiring that an investigator provide only the input parameters and data (Fig. 1). When an analysis is launched, it is executed on the specified remote server, with the job execution status displayed within the cell. When the job completes, links to the result files are presented in the notebook cell and can be used easily as input to further analyses (Fig. S1). Thus, to the user, the entire analysis workflow appears to run seamlessly within the notebook.

**Fig. 1.**
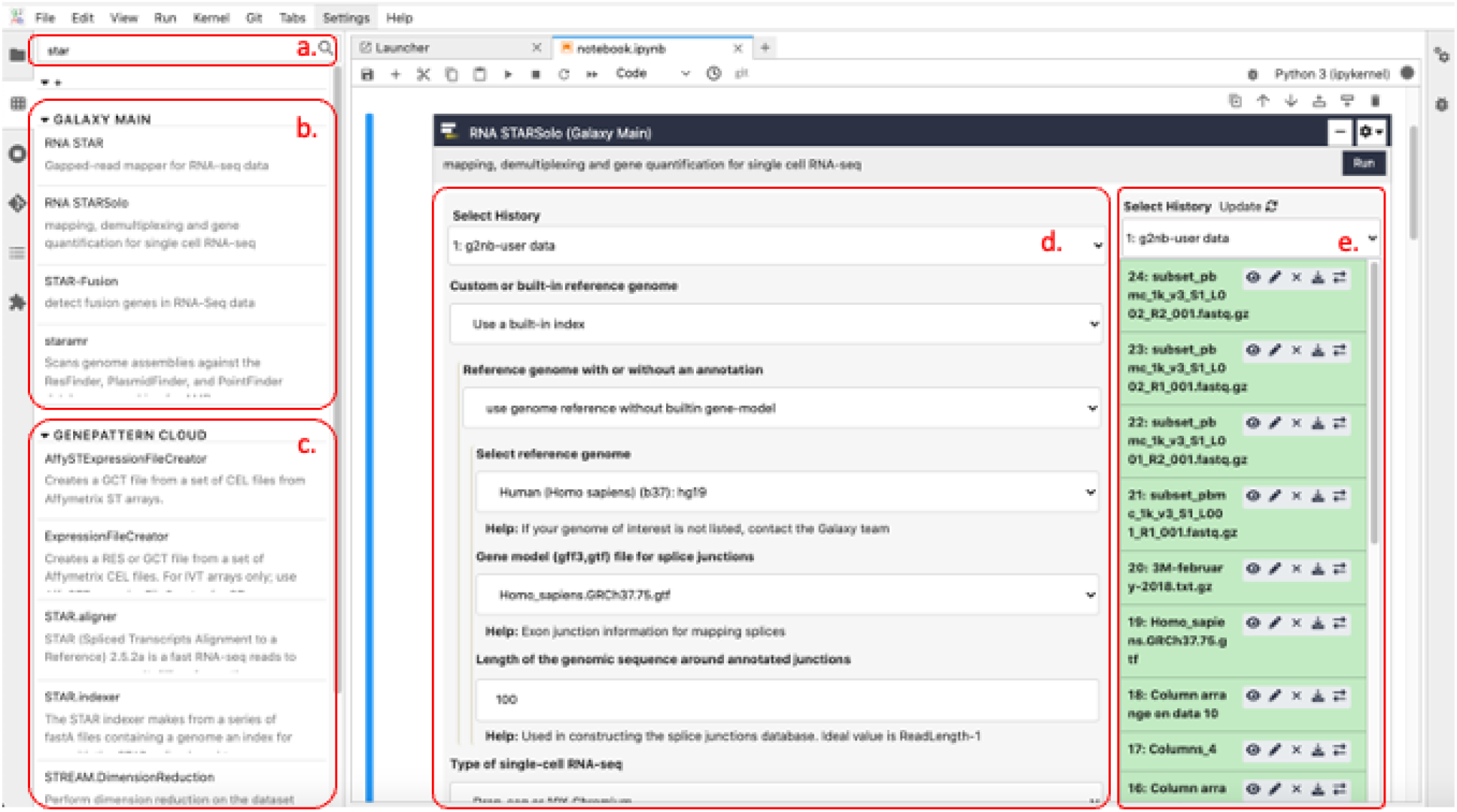
g2nb user interface. A tools panel (a-c) allows users to search for and select from any server that the user has logged into. In this example the user (a) searches for the STAR sequence alignment tool. Tools with STAR in their name or description are displayed for the two servers currently connected: (b) the Galaxy Main server and (c) the GenePattern cloud server. (d) A Galaxy analysis cell shows the interface to the STARsolo tool on Galaxy Main after it has been selected by the g2nb user. (e) input files are selected as they would in the original platforms. Here a Galaxy history is displayed, with possible user input files. Figs. S1-S3 show GenePattern, IGV, and Cytoscape in their g2nb analysis cell formats.

The g2nb environment provides capabilities for interactively visualizing various genomics datatypes by including the JavaScript versions of Cytoscape[5] and IGV[6], as well as standard heatmap, dendrogram, and other plots provided in GenePattern and Galaxy. Visualizations appear within notebook cells and also can be launched in independent windows if more space is required. For the visualization tools, g2nb takes advantage of functionality already provided by those projects to run within the JupyterLab environment. The Cytoscape project has released CyJupyter (https://github.com/cytoscape/cytoscape-jupyter-widget), a visualizer that supports a growing subset of the capabilities of the standard desktop Cytoscape application. Similarly, the developers of the Integrative Genomics Viewer (IGV) have released igv-notebook (https://github.com/igvteam/igv-notebook), which packages the igv.js (https://github.com/igvteam/igv.js) JavaScript IGV browser for use in notebook environments. While these tools typically require Python knowledge to install and run in the notebook environment, g2nb makes them available as launchable tools within its tool panel (Figs. S2, S3).

To facilitate the seamless flow of data into a notebook and through notebook cells, the g2nb environment provides several new JupyterLab enhancements. These include: presenting Galaxy histories and GenePattern result files within notebook cells, thereby allowing users to easily select files from previous analyses; displaying a list of all analyses in a notebook that can receive a particular result file as input; and running a sequence of cells as an end-to-end workflow, including cells that launch jobs outside the notebook on Galaxy or GenePattern servers. Because links to result files within analysis cells reference remote Galaxy or GenePattern servers, any necessary data transfers are handled automatically. We have also incorporated the Globus[7] file transfer protocol into the g2nb interface, allowing users to robustly transfer files of any size between the g2nb workspace and any Globus endpoint, including the user’s local storage. For programmers, we have implemented features to facilitate transfer between Python objects and Galaxy and GenePattern jobs. In a g2nb notebook, a Python variable can be provided as an input parameter to Galaxy and GenePattern analyses. The g2nb environment will evaluate the variable and pass its value to the analysis. Additionally, rather than having to download result files and read them into Python objects, users can automatically load the contents of a result file directly into a Python variable or, for array-type data, into a Pandas dataframe.

For notebook authors, g2nb provides a *User Interface Builder* that allows them to present Python code cells in a more user-friendly format. Notebooks frequently include cells with large blocks of code that require no interaction other than being run by the user, or for the user to provide a small amount of information. The User Interface Builder allows authors to create a web form interface to a cell, exposing only those parameters they wish users to enter. Notebook authors can specify data inputs, text or numeric entry or dropdown lists, and can tailor the interface in many ways to the specifics of their code. The underlying code is always available via a toggle button.

We provide a freely available online workspace, g2nb.org, where investigators can create and run notebooks, share them with collaborators, and publish them for general use. The workspace includes several components to help scientists use the environment: a growing library of g2nb notebooks that implement common analysis workflows that investigators can copy and use as templates for their own research. Notebooks in the collection currently support several ‘omics modalities. Single-cell analysis, including preprocessing, clustering, cluster harmonization, pseudotime trajectory analysis, and RNA velocity analysis, are available via notebooks that incorporate the Seurat[8], Conos[9], STREAM[10], and scVelo[11] tools; a number of gene set enrichment analysis modalities including “classic” two-phenotype GSEA[12], single-sample GSEA[13]; and general machine learning methods for clustering, classification, and dimension reduction. To address the frequent problem of notebooks having incompatible dependencies, g2nb provides *project spaces*. Each project space provides a separate context that contains its own notebooks, packages, libraries, and files. Investigators can share projects with collaborators and publish projects on the g2nb workspace.

While several projects have attempted to combine notebook systems with bioinformatics platforms, g2nb is novel in that it incorporates multiple platforms within the JupyterLab interface, with functionality to seamlessly transfer data between tools. The Galaxy project previously integrated Jupyter as a tool within its web interface, allowing users to run a Jupyter notebook as a step in their analyses [14]. This approach complements the one in g2nb, where the notebook provides the user environment and Galaxy tools can be launched from within it in combination with other software packages. Many other tools and resources provide packages that allow them to be used in a Jupyter-based environment, but these are generally code libraries that do not make use of the notebook’s features beyond their use in code cells.

## Code Availability

The g2nb platform is available for use at g2nb.org. For those who wish to run or host the g2nb environment on a local system, it is available as the g2nb/lab and g2nb/workspace Docker containers at hub.docker.com. The source code for the g2nb architecture is available at github.com/g2nb under a BSD-style open source license.

## Acknowledgements

The work was funded by NIH grants U24CA194107 to JPM and NIH NHGRI U24HG006620 and NIH NCI U24CA231877 to DB.

. We thank Jim Robinson and Dexter Pratt for helpful discussions.

