## Supplemental Data for "Genomics to Notebook (g2nb): extending the electronic notebook to address the challenges of bioinformatics analysis"

**Figure S1. Features for working with result files**. A g2nb notebook is displayed with three GenePattern cells. The result file from the ssGSEA analysis (a) displays a context menu when clicked, showing (b) standard options for displaying and downloading the file, as well as for automatically loading the files contents into a Pandas dataframe (this option is available for files containing data in an array format) and a Python variable (Send to Code). The menu also presents a list of the analysis cells and their inputs in the notebook that can accept that file (c). Here, the results of ssGSEA can be used as input to the HierarchicalClustering analysis cell or the HeatMapViewer visualization cell. When one of these is selected, the file appears as the input of the corresponding analysis. In each case (b-c), the result data is automatically downloaded and transferred to the appropriate destination, providing a seamless user experience.

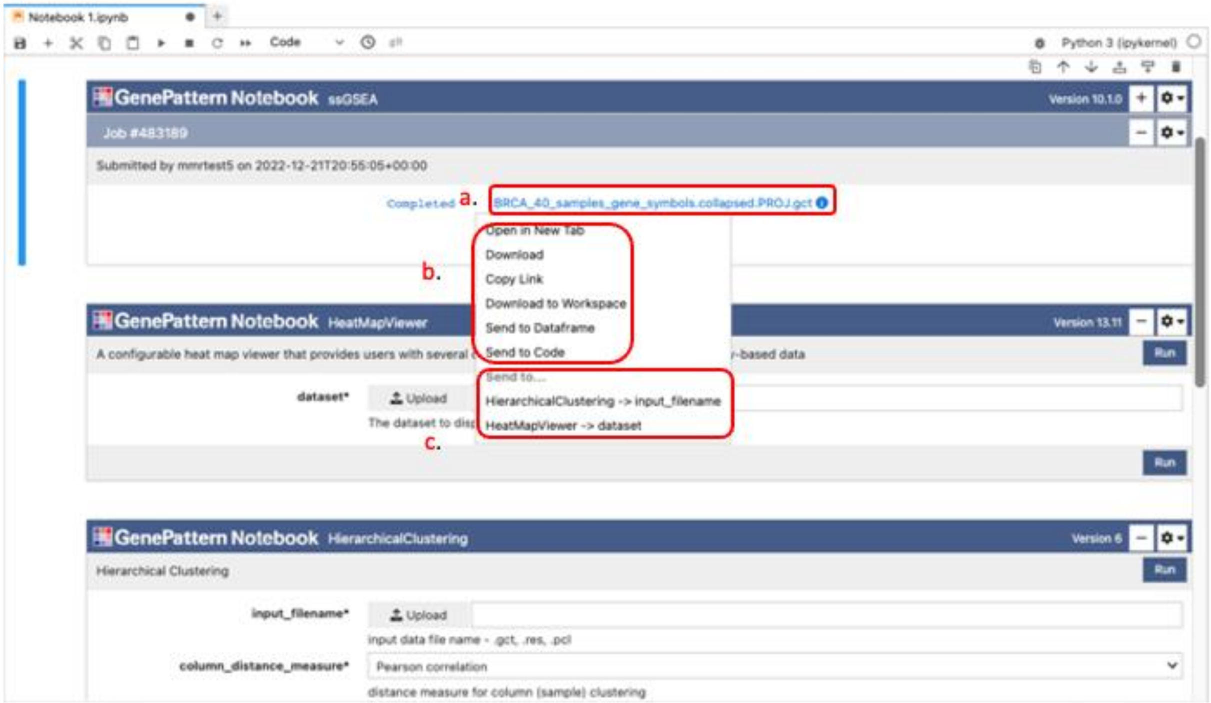

**Figure S2. IGV in g2nb**. To insert an IGV cell into a g2nb notebook, a user (a) selects the Integrative Genomics Viewer tool in the toolbar. After the user specifies the input data, (b) the IGV viewer is displayed as a notebook cell, with all of IGV's interactive capabilities, including zooming, panning, multi-locus view, searching by gene name or genomic locus, etc. Here IGV is displaying aligned reads from whole genome sequencing, with the view zoomed in to gene GSTT1. The green color represents mismatches between the data and the reference genome, clearly indicating a SNP in the sequenced sample.

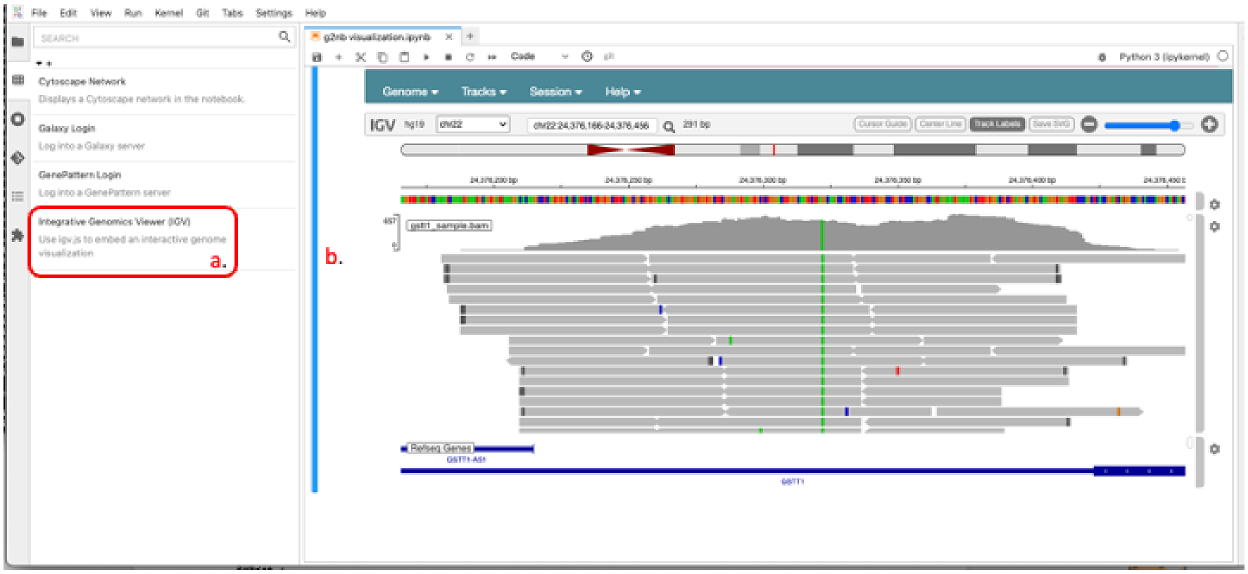

**Figure S3. Cytoscape interface in g2nb**. The g2nb environment uses the JavaScript-based Cy-JupyterLab package to implement a subset of available Cytoscape functionality. The user (a) selects the Cytoscape tool in the toolbar. After the user specifies the input data, the Cy-JupyterLab interface (b) is displayed as a notebook cell, with capabilities including zooming, panning, manipulation of nodes, and other functionality as described in the GitHub repository at https://github.com/cytoscape/cy-jupyterlab. Here the Cytoscape tool is displaying the MTOR signaling pathway as retrieved from the NDEx[18] resource (https://www.ndexbio.org/viewer/networks/34540ca5-1e5f-11e8-b939-0ac135e8bacf)

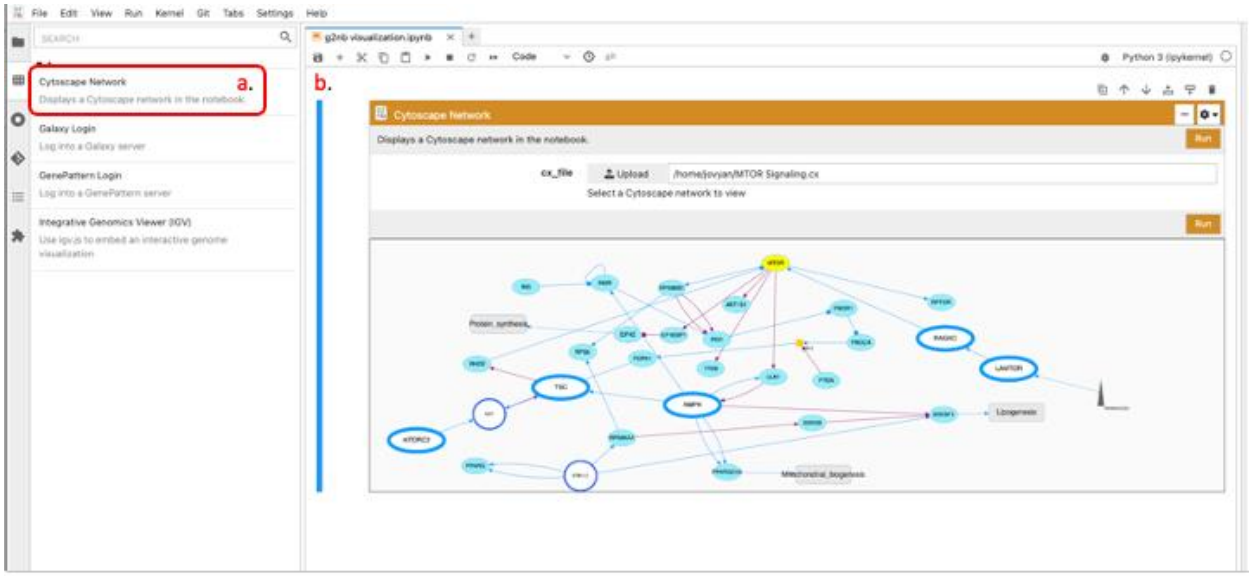
